# Mechanics of knee meniscus results from precise balance between material microstructure and synovial fluid viscosity

**DOI:** 10.1101/2024.05.15.594315

**Authors:** Camilo A.S. Afanador, Stéphane Urcun, Ivo F. Sbalzarini, Stéphane P.A. Bordas, Olga Barrera, Mohammad Mahdi Rajabi, Romain Seil, Anas Obeidat

## Abstract

The meniscus plays a crucial role in the biomechanics of the knee, serving as load transmitter, and reducing friction between joints. Understanding the biomechanics of the meniscus is essential to effective treatments of knee injuries and degenerative conditions. In this study, we used two central meniscus samples extracted from a human knee and acquired high-resolution ***µ***-CT images. Using an implicit immersed boundary technique, we reconstructed two 3D computational models of the menisci. By eroding the channels of the original meniscus geometry, we created new microstructures with varying porosities (**0.53** to **0.8**) whilst preserving the connectivity of the porous structure. We investigate the fluid dynamics of the meniscus using a mesh-free numerical method, considering various inlet pressure conditions and analysing the fluid flow within the microstructures. The results of the original microstructure associated with a physiological dynamic viscosity of synovial fluid are in accordance with biophysical experiments on menisci. Furthermore, the eroded microstructure with a **33%** increase in porosity exhibited a remarkable **120%** increase in flow velocity. This emphasises the sensitivity of meniscus physiology to porous microstructure properties, showing that detailed computational models can explore physiological and pathological conditions, advancing further knee biomechanics research.

## 1 Introduction

The biomechanical functions of menisci in the knee include load transmission, friction reduction between joints [1–3], and possibly shock absorption. The shock-absorbing function of the meniscus, while commonly cited, is however a subject of ongoing debate [4]. Prior to 1980, menisci were often considered superfluous and were frequently removed after injury, leading to negative consequences such as arthrosis in patients [3]. The understanding of meniscus function has improved significantly, with multiple aspects believed to contribute to their energy dissipation ability, including the geometry and the biochemical composition of the microstructure [1–3, 5–10].

In this study, menisci are considered as a porous medium. This porous medium contains a structural solid scaffold mainly composed of type I collagen (75%), large hydrophilic molecules termed proteoglycans (6%) [5]. This solid scaffold is saturated by synovial fluid, the dynamic viscosity of which depends on the meniscus’ health and its physiological conditions [11].

The main component of the structural scaffold, the collagen fibers which ensure its mechanical integrity [12], contributes to energy dissipation by its structure ensuring a circumferential tensile stress [6, 7]. The chemical composition of the structural scaffold, particularly the presence of proteoglycans, plays a crucial role in defining the shock-absorption properties [5]. Proteoglycan are large hydrophilic molecules that ensure tissue elasticity under small loads [5]. Under larger loads (> 100 kPa), the interstitial fluid starts to flow, causing menisci to behave as a viscous porous medium [8].

We only found few quantitative studies that used numerical simulations to relate the mechanical and structural properties of the meniscus [13–15]. These studies have provided insights into the viscoelastic properties and the anisotropy of permeability due to collagen fiber orientation in the meniscus [15, 16].

Advances in computing power and microscale characterisation techniques, such as micro-Computed Tomography scanners (*µ*-CT), magnetic resonance velocimetry (MRV), and X-ray microtomography (XMT), have enabled the study of fluid flow and transport processes at the microscopic level in porous media [17–20]. These techniques allow for accurate reconstruction of the pore structure of porous media in three dimensions, which can then be used as input for computational fluid dynamic (CFD) simulations at the pore scale [21, 22]. Here we use these computational advances to disentangle the contributions of chemical, structural, and geometric factors to the energy dissipation ability of human menisci, building on existing literature and research findings [1–3, 5–10].

We consider two healthy meniscus samples from a human knee [10] imaged by high-resolution *µ*-CT (6.25 *µ*m resolution). For these samples, we reconstructed a 3D computational geometry model using an implicit immersed boundary technique. We then solve an alternative formulation of the incompressible Navier-Stokes equations (the Entropically Damped Artificial Compressibility equations) inside the resulting microstructures, using different inlet pressure boundary conditions representative of different loading scenarios.

We find that by surpassing a certain pressure threshold, the flow in the meniscus qualitatively changes from a slow creeping flow to a pressure-dominated laminar flow. The existence of these two qualitatively different flow regimes is typically associated with healthy menisci. Increasing the porosity of the meniscus microstructure enhances hydration, but excessive porosity leads to pathological responses, interfering with joint function, affecting tissue nutrient exchange, and leading to joint degeneration [23–27]. We elucidate the mechanism behind this by eroding the meniscus geometry to construct microstructures ranging from a porosity of 0.53 to a porosity of 0.80 at constant connectivity of the pore structure. Of course, an artificially increased porosity as high as 0.8 is not physiological. However, the purpose of dilating the microstructure to such limits is to extrapolate the impact of the meniscus’ microstructural characteristics on its macroscopic properties. We find that eroded geometries: (i) no longer exhibit the two flow regimes and (ii) display a non-linear relationship between porosity and velocity magnitude. For instance, a 30% increase in porosity leads to a 120% increase in velocity velocity.

Taken together, the fully resolved pore-scale computer models of meniscus geometry and direct numerical fluid flow simulations within them allow us to accurately capture the geometry–mechanics trade-off in menisci. On the one hand, fluid flow in a meniscus should be sufficiently slow and dampened for it to exhibit shock absorbing properties. On the other hand, though, some flow must exist in order to transport metabolites and nutrients in and out of the tissue. This requires a fine-tuned balance between the geometric microstructure of the meniscus and the viscosity of the synovial fluid. Using the present simulations, we were able to understand this balance mechanistically and predict impact of meniscus microstructure on mechanical function under fully controlled conditions.

## 2 Results

In Fig. 1 we outline the main steps of this work. In the study by Agustoni *et al*. [10], two meniscus samples **S1G0** and **S2G0** were extracted. In order to obtain better imaging contrast, the samples were freeze-dried prior to the *µ*-CT (6.25*µ*m) scans and therefore contained only two phases, solid and void. It has been previously shown [10] that the specific freeze-dry procedure preserves the range of the pore sizes,in comparison to the results obtained by Vetri et al. [28] using confocal multi-photon microscopy. We can not exclude the existence of an artificial porosity provoked by this experimental process, as we acknowledge that no observation technique is without limitations. Then, the *µ*-CT scans were used to produce the 3D volume STL (Standard Triangle Language) format geometry of the meniscus samples, which serves as an input to our work, Fig. 1(A). The ex-vivo sample **S1G0** had a diameter of 3.5 mm and a length of 4.6 mm. The produced STL had 5.68 million vertices and 11.55 million faces. The ex-vivo sample **S2G0** had a diameter of 1 mm and a length of 3.13 mm. The produced STL geometry had 3.55 million vertices and 7.14 million faces, (see From sample to computational domain). A noteworthy emphasis is placed on the distinct size different between the two samples, with **S1G0** presenting a diameter that is 3.5 times greater than its counterpart, **S2G0**.

**Fig. 1:**
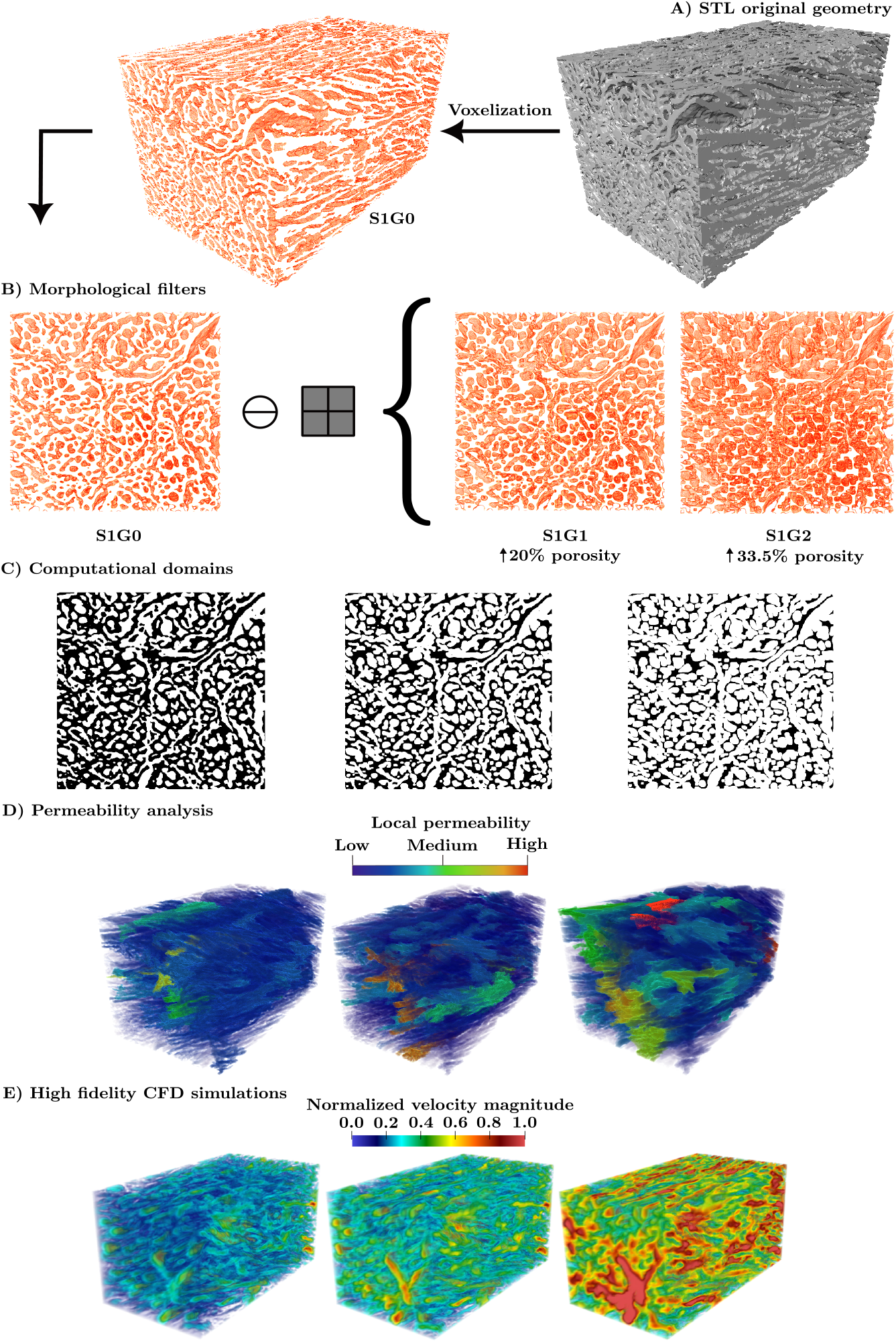
Workflow outline for investigating the effect of microstructural properties on fluid flow in human meniscus: **(A)** The 3D STL geometry of the microstructure volume of the human meniscus (**S1G0**) is constructed from high-resolution *µ*-CT (6.25*µ*m). **(B)** The meniscus channels are eroded by applying morphological filters (erosion) to obtain two additional geometries (**S1G1** and **S1G2**) with different microstructural properties. This controlled erosion filter changes the microstructural properties of the meniscus, mainly the porosity. The resulting geometries allow us to study the effect of the meniscus’s wall degradation on its5 functionality, as shown in Table 1. **(C)** The computational domain of the different meniscus geometries (**S1G0, S1G1**, and **S1G2**) with implicit boundary representation is created using an in-the Brinkman penalisation. (**D**) Flux analysis at the pore scale for the three structures yields a distribution of local permeability, Local permeability distribution is characterised as volume regions of the porous network with diameters of channel exceeding 4 voxels. Blue regions indicate areas of lower local permeability and correspond to smaller volume magnitudes. Conversely, red regions represent higher local permeability, signifying areas with greater volume magnitudes. (**E)** High-fidelity computational fluid dynamics (CFD) flow simulations are conducted using the Discretisation-Corrected Particle Strength Exchange (DC-PSE) method. Evolution of the velocity field with respect to the porosity: from **S1G0** to **S1G1** and **S1G2**, the porosity increases by 20.5% and 33%, respectively, and the maximal velocity increases by 30% and 120%, respectively.

There is a clinical need to study the effect of the meniscus wall degradation on its functionality. Meniscus degeneration, however, is a multi-faceted process that encompasses not just geometric alterations to the microstructure, but also changes in the biochemical composition, material properties, and structural organisation of the meniscus. Understanding meniscus degeneration will therefore ultimately require a holistic approach that goes beyond morphological alterations [29, 30]. As a first step, we here focus on the effect of microstructure geometry on fluid mechanics in the meniscus, which our computational model enables us to study in isolation. For this, we apply a uniform erosion morphological filter with two different characteristic lengths to **S1G0**, modifying its porosity to obtain two new “degraded” meniscus geometries (**S1G1** and **S1G2**) while preserving the connectivity of the porous network, Fig. 1(B). The 3D volume of the three geometries is then represented in the mesh-free computational domain as particles (point clouds). Due to the complexity of the meniscus geometry, the Brinkman penalisation technique is applied for implicit representation of the complex boundaries [31], Fig. 1(C). In Fig. 1(D) the local permeability distribution of the three porous geometries is presented. This distribution allows the identification of areas in which the flow will tend to move more easily (high local permeability), see Fig. 1(D) and inline with our numerical results in Fig. 1(E), further validating the significance of local permeability in influencing fluid flow behavior within the porous medium. The Entropically Damped Artificial Compressibility (EDAC) solver for The Discretisation-Corrected Particle Strength Exchange (DC-PSE) [32] is utilised to model the three dimensional viscous fluid flow inside the meniscus by solving the EDAC formulation, in Fig. 1(E). Finally, we examine the the impact of inlet pressure and dynamic viscosity on fluid flow, considering various porosity values across different microstructures, Figs. 4 and 5.

**Table 1:** Computed volume fraction v_*f*_ and microstructure statistics for the simulation domains.

| Geometry | erosion length of the structural element $\ell_c$ | Average porosity | Non-connected clusters | $v_f$ of connected voids | Average tortuosity |
| --- | --- | --- | --- | --- | --- |
| S1G0 | 0 | 0.534462 | 247 | 0.997377 | 2.4 |
| S1G1 | 2 | 0.672338 | 180 | 0.999631 | 1.7 |
| S1G2 | 4 | 0.803577 | 69 | 0.999928 | 1.4 |
| S2G0 | 0 | 0.510832 | 92 | 0.998975 | 2.2 |

### 2.1 Geometrical parameter analysis

Before analysing three crucial geometrical parameters, namely porosity, average tortuosity, and connectivity, it is necessary to examine how the microstructural properties of the meniscus impact its functionality.

We modify the healthy sample **S1G0** outside its physiological configurations, the sample undergoes a morphological filter (erosion filter) to artificially increase the porosity by increasing the channel diameters. This filter has a significant impact on the properties of the porous medium, see Table 1. In the table, 𝓁_*c*_ represents the structural element length, which corresponds to the size of the voxel being eroded. Applying the erosion filter of length (structural element 𝓁_*c*_) 2 and 4 voxels, the porosity of **S1G0** is artificially increased by 20.5% and 33.5%, respectively. This leads to the creation of two new meniscus geometries, **S1G1** and **S1G2**, as presented in Fig. 1(B,C). Tortuosity provides a measure of the difficulty level for fluid to flow within a porous network. When the length scale of the erosion filter is increased, the average tortuosity decreases, indicating that the porous network becomes more permeable.

The non-connected clusters in a porous medium refer to isolated regions within the sample where there is no possibility of fluid flow due to a lack of connectivity to other regions of the sample. In other words, these clusters represent areas where fluid flow is hindered or impossible. The numbers provided in Table 1, in the volume fraction v_*f*_ of connected voids column, indicate that the porous network in the original geometries **S1G0** and **S2G0** have a high degree of connectivity, > 99.7% of the volume fraction. This is important for a living tissue, as it ensures that there are no “dead” regions without fluid exchange. This high degree of connectivity is preserved also in the eroded geometries, showing that the sample porosity can be modified without significantly altering the overall connectivity of the porous network (note that for **S2G0**, the number of non-connected clusters is representative of a smaller volume compared to **S1G0**), see Table 1.

The volume fraction in relation to the normalised pore diameters is shown in Fig. 2(B), which provides insight into the homogeneity or heterogeneity of the pore structure. The wide distribution of pore diameters in **S1G0** and **S2G0** indicates a high level of heterogeneity, while the peak at 0.9 of the maximum diameter in **S1G2** indicates a high level of homogeneity.

**Fig. 2:**
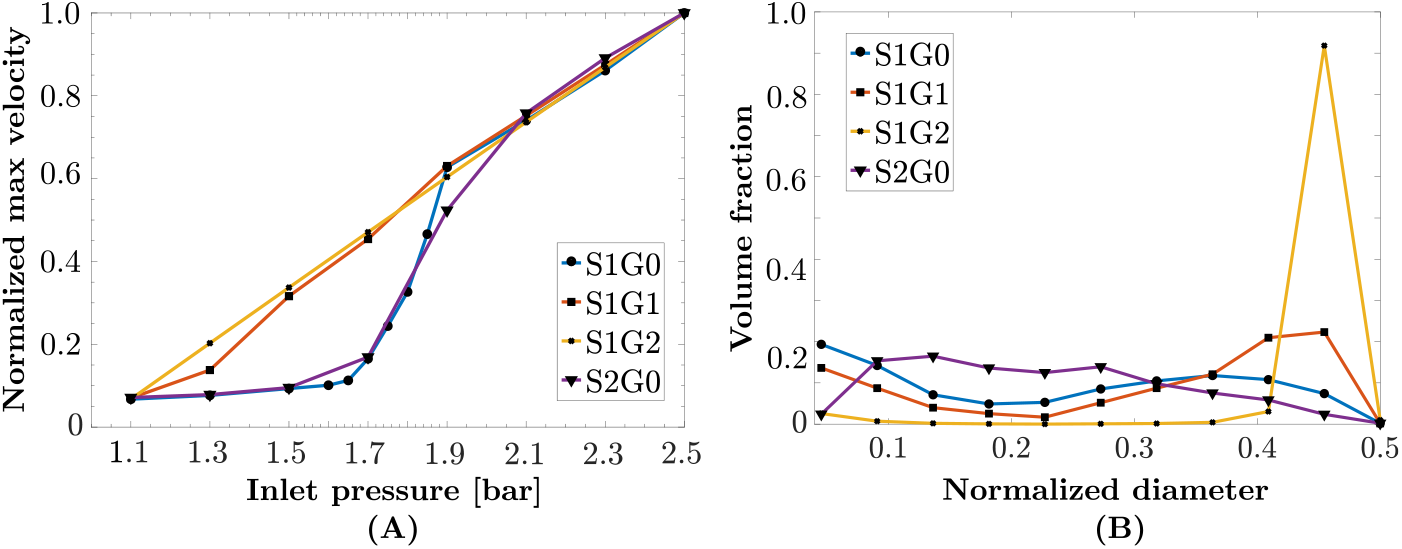
**(A)** The Normalised maximum flow velocity vs. inlet pressure; original geometry **S1G0** (•, blue); eroded geometry **S1G1** (◼, red); eroded geometry **S1G2** (×, yellow); original geometries 2 **S2G0** (▼, purple). **S1G0** and **S2G0** show a threshold behavior with respect to the inlet pressure, in accordance with the flow regimes described by Fithian *et al*. [6]. The eroded geometries **S1G1** and **S1G2** lose this property, but keeping the same behavior at high pressures. **(B)** The volume fraction occupied by different normalised pore diameters. The wide distribution of pore diameters in **S1G0** (•, blue) and **S2G0** (▼, purple) indicates a high level of heterogeneity, while the peak at 0.9 in **S1G2** (×, yellow) indicates a relatively homogeneous geometry.

The histograms and fitted probability density functions for porosity, along with the average tortuosity, are presented for the four geometries in Fig. 3. The histograms provide insights into the distribution of porosity and tortuosity within each microstructure, revealing unique characteristics and variations among the studied microstructures.

**Fig. 3:**
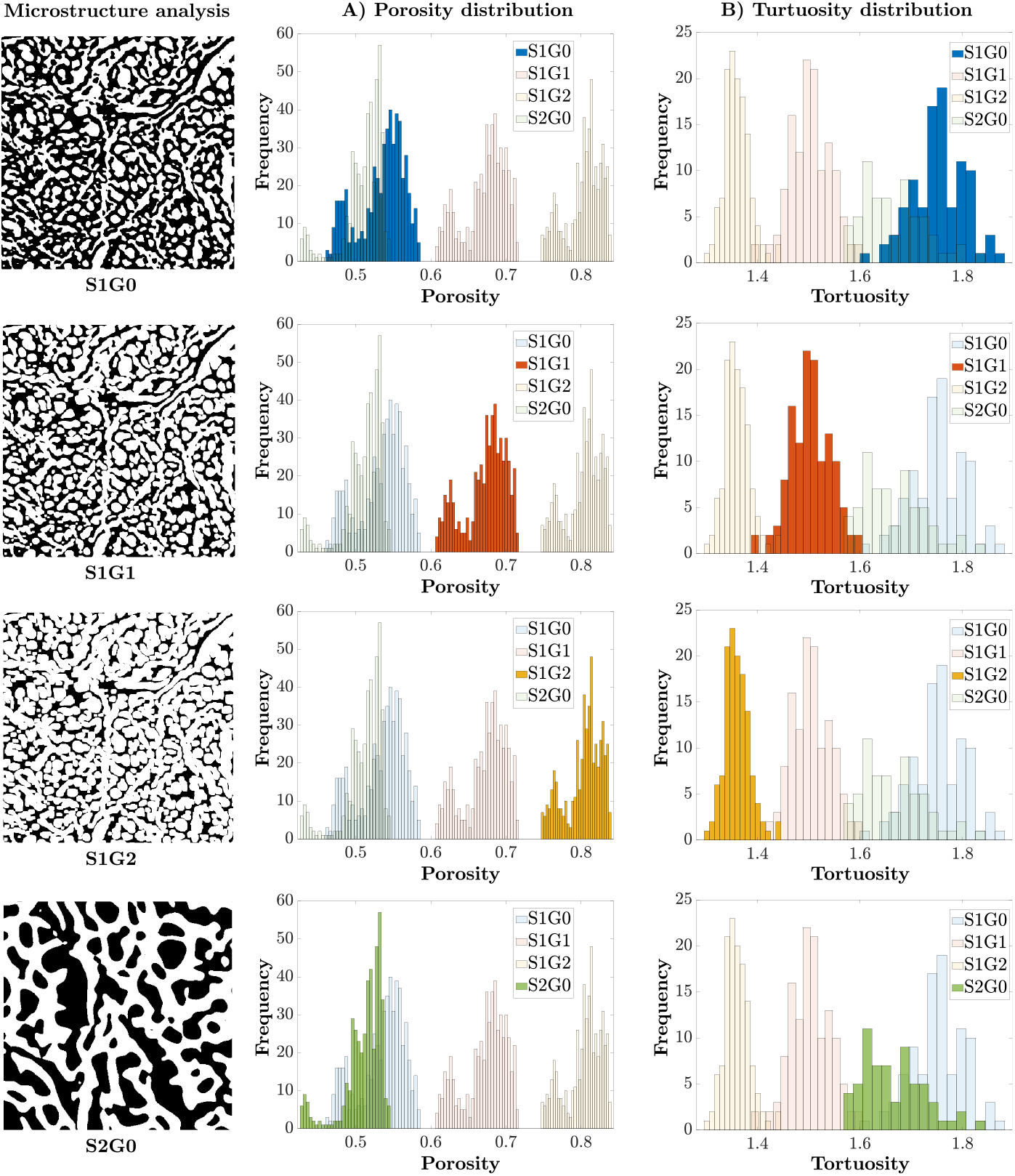
Comparative analysis of four distinct microstructure:**S1G0, S1G1, S1G2** and **S2G0**. Each row represents one microstructure commencing with a visual representation followed by it respective porosity and tortuosity. For a better visualisation, we note that the corresponded histograms to the the sample are in full colour, while the distributions for other samples are rendered transparently. Additionally, the microstructure of S2G0 is magnified 3.5 times to ensure consistent representation with the other samples.

We note that the porosity of the original geometry (53±5%), obtained by *µ*CT scans analysis, is within the range of the current literature (from 34.1% [33] to 65% [34]). We also note that the mean value of tortuosity in the original geometries is 2.3, close to the value of 2, commonly used in the porous medium literature for living tissues [35–37]. As the porosity increases in **S1G1** and **S1G2** the tortuosity decreases, indicating a more permeable porous medium.

The porosity values of **S1G0** and **S2G0** (0.53 ± 0.05) are in accordance with the porosity measurements by *µ*CT-scans reported in the literature. We note a degradation of porosity in **S1G1** and **S1G2** as a result of applying the morphological filter. The mean value of tortuosity is close to 2 for **S1G0**, and **S2G0**, which is the common value used the in porous medium literature for living tissues [35–37]. As the porosity increases in **S1G1** and **S1G2** the tortuosity decreases, indicating more permeable porous medium.

### 2.2 Effect of inlet pressure on the flow regime

The focus of our study is to investigate how fluid flow is affected by varying pressure at the inlet, with different microstructural properties (healthy and pathological menisci). To accomplish this, we apply initial pressure boundary conditions at the inflow using a range of values from 1.1 to 2.5 bar (see Simulation setup, initial condition and boundaries). According to Fithian *et al*. [6], synovial fluid can exhibit two types of flow behavior: (1) visco-elastic behavior, which is mediated by proteoglycans of the extra-cellular matrix, (2) porous-permeable behavior, which occurs when the synovial fluid is subjected to flow through a hydraulic pressure gradient or matrix compaction.

The results of our numerical simulations are summarised in Fig. 2(A), where those two regimes of the velocity with respect to inlet pressure are clearly seen for the original geometries **S1G0** and **S2G0**. It is clear that only the original, healthy geometries **S1G0** and **S2G0** exhibit two flow regimes with a threshold of around 1.7bar inlet pressure. This threshold behavior is not seen in any of the eroded geometries **S1G1** and **S1G2**, where porosity no longer limits fluid flow.

In healthy samples (**S1G0** and **S2G0)** this surprising qualitative change in behavior is not an artifact of the mathematical flow model used (see Mathematical modelling), as the pressure-velocity relationship is locally linear, but an intrinsic geometrical property of the original sample itself and the physiological viscosity of the synovial fluid. For a better understanding of this behavior, the histograms of velocities with an inlet pressure of 1.7 and 2.1 bar for **S1G0** are provided in Fig. 4(B-1). The simulated flow for sub-critical inlet pressure (blue in Fig. 4), presents a general trend toward extremely small velocities (visco-elastic). Specifically, in **S1G0**, more than 50% of the porosity contains flow below 5% of maximal velocity (1.53·10^*−*3^ m/s), and this trend is even more obvious for **S2G0** (Fig. 4(B-2)). Above the threshold of 1.7 bar, (red in Fig. 4), the model predicts a porous-permeable flow behavior, which means that the flow is dominated by the pressure gradient. The velocity distribution becomes more Gaussian in shape and centered around about 33% of the maximal velocity. This is a clear indication that the flow behavior has transitioned to the porous-permeable regime.

**Fig. 4:**
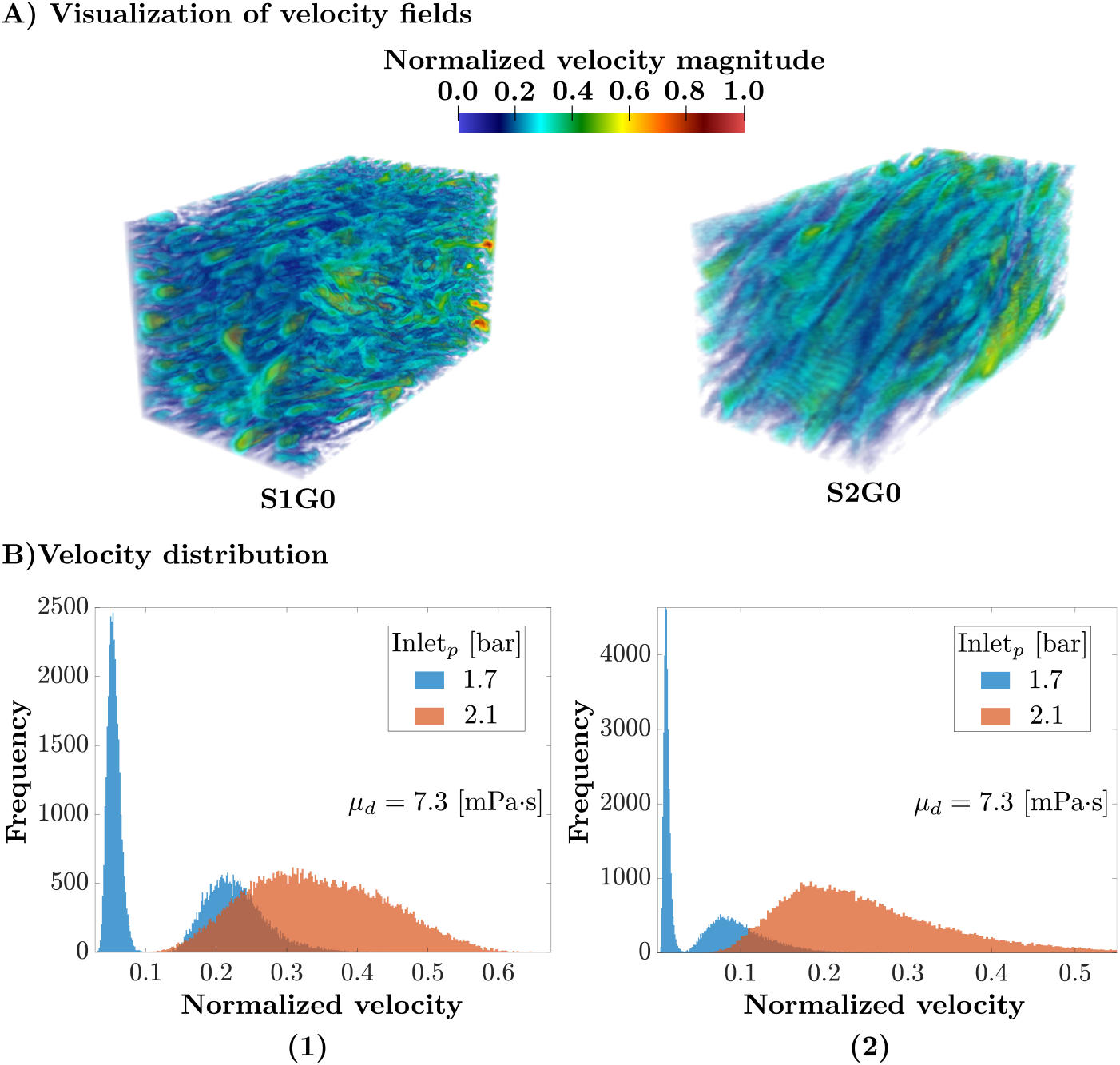
**(A)** visualisation of the normalised (to the maximum value in S1G0) flow velocity fields for **S1G0** and **S2G0**, with inlet pressure 2.1 bar. **(B)** flow velocity distribution histograms; in blue, inlet pressure 1.7 bar; in red, 2.1 bar. The velocity is normalised with the maximum velocity observed for inlet pressure P_inlet_ = 2.1 bar, with the reference viscosity *µ* = 7.3·10^*−*3^ Pa·s. **(B-1)** The histogram of the non-zero velocity field for **S1G0**. The flow presents a general trend toward extremely small velocities (visco-elastic), more than 50% of the porosity contains flow below 5% of maximal velocity (1.53·10^*−*3^ m/s). The synovial fluid transitions to porous-permeable behavior, which means that the flow is dominated by the pressure gradient, at super-critical pressure. The velocity distribution then becomes more Gaussian in shape and centered at around 33% of the maximal velocity.**(B-2)**The same histograms for sample **S2G0**, showing the threshold behavior even more clearly.

The synovial dynamic viscosity has an impact on the mechanical function of the meniscus. A viscosity below the physiological range is for example observed in cases of aseptic loosening after total joint replacement [11]. This change in viscosity affects the flow behavior of the synovial fluid, as evidenced by the loss of the threshold behavior observed under healthy conditions, see Fig. 5(C-1). This confirms that the dual flow behavior is not solely influenced by the geometry of the samples, but also by the physiological properties of the synovial fluid.

**Fig. 5:**
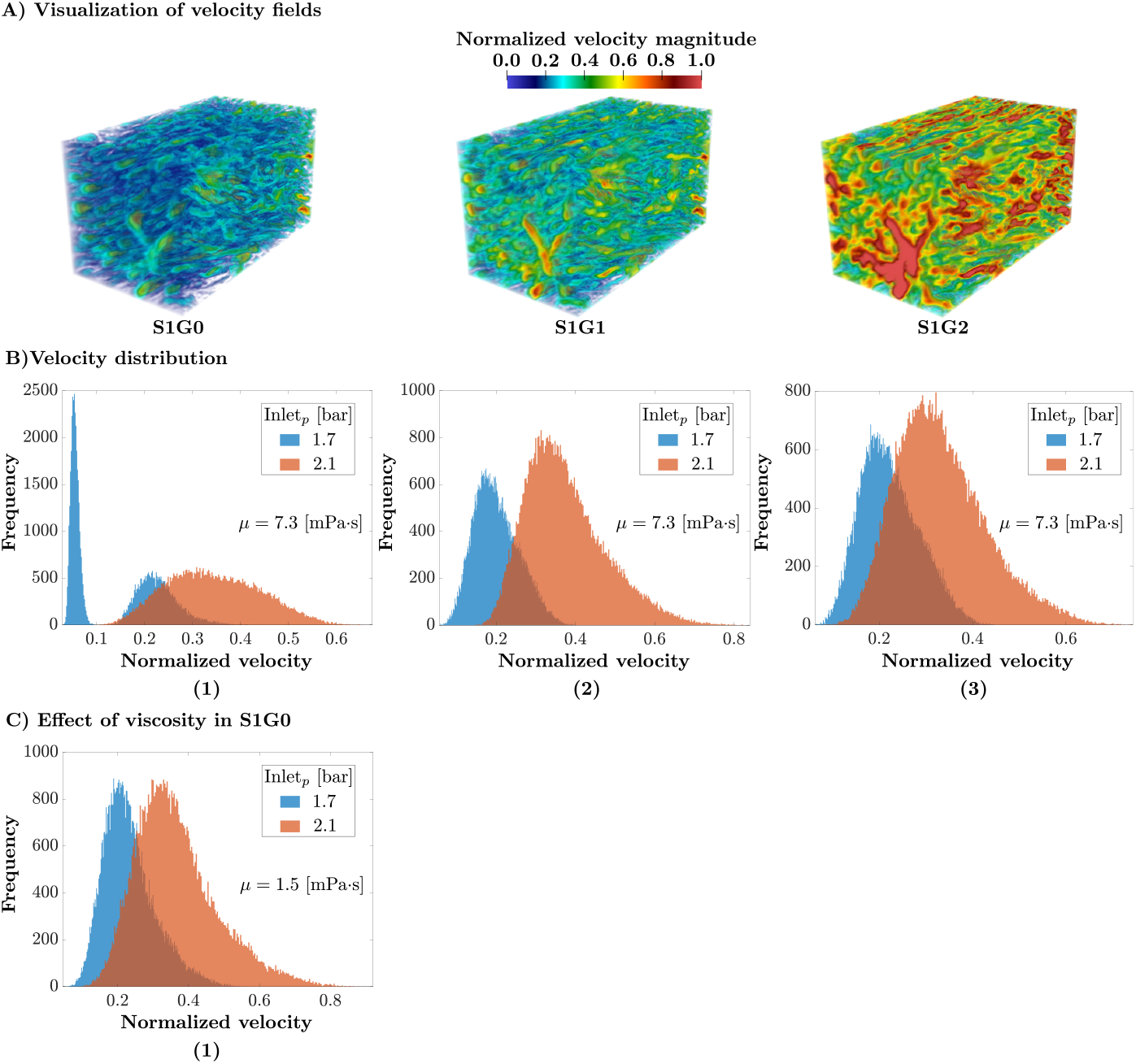
**(A)** visualisation of the normalised (to the maximum value in S1G0) flow velocity fields for **S1G0, S1G1** and **S1G2**, with inlet pressure 2.1 bar. The normalised velocity in **S1G2** is 120% higher than in **S1G0**, whereas it is only 30% higher in **S1G1**, highlighting the nonlinear effect of pore size on flow dynamics in the meniscus. **(B)** flow velocity distribution histograms; in blue, inlet pressure 1.7 bar; in red, 2.1 bar. The velocity is normalised with the maximum velocity observed for inlet pressure P_inlet_ = 2.1 bar, with the reference viscosity *µ* = 7.3·10^*−*3^ Pa·s. **(B-1)** The flow velocity distribution histogram of the non-zero velocity field for **S1G0**. The flow presents a general trend toward extremely small velocities (visco-elastic), more than 50% of the porosity contains flow below 5% of maximal velocity (1.53·10^*−*3^ m/s). The synovial fluid transitions to porous-permeable behavior, which means that the flow is dominated by the pressure gradient, at super-critical pressure. The velocity distribution then becomes more Gaussian in shape and centered at around 33% of the maximal velocity. **(B-2)** The histogram of the non-zero velocity field for sample **S1G1. (B-3)** The histogram of the non-zero velocity field for sample **S1G2**. The results of the two previous structures suggest that the geometrical properties of the meniscus significantly affect the behavior of fluid under varying pressure. Here, the threshold behavior is lost in both samples with flow always in the porous-permeable regime. **(C-1)** The flow velocity histograms for **S1G0** with a lower synovial fluid viscosity *µ* = 1.5·10^*−*3^ Pa·s, characteristic of aseptic loosening after total joint replacement [11], where the dual behavior is lost. This confi12rms that the two flow regimes observed in the healthy samples are not solely due to the geometry of the samples, but also depend on the physiological properties of the synovial fluid, specifically as its viscosity.

The flow transition also disappears upon meniscus degeneration, modeled by an increased porosity (+20.5% for **S1G1**, accompanied by a −29.1% decrease in tortuosity), see Fig. 5(B-2). The meniscus then shows a linear flow behavior with respect to the inlet pressure. The same is observed in **S1G2** (porosity increases of +33.5% and tortuosity decreases of −41.6%), see Fig. 5(B-3). Since no other simulation parameter changed, this qualitative change can be attributed purely to intrinsic geometrical properties of the samples at physiological values of synovial fluid viscosity. These results are summarised Fig. 2(A), where the threshold behavior of velocity with respect to inlet pressure is only visible for the original healthy geometries (**S1G0** and **S2G0**).

### 2.3 The effect of the pore size on the maximum velocity

We perform numerical simulations of the fluid dynamics of the synovial fluid inside the fully-resolved porous microstructure geometries of two human meniscus samples **S1G0** and **S2G0**. The details of the simulation method are given in the Materials and Methods section and have been verified and validated elsewhere [32].

In order to see how the average pore diameter influences the fluid flow velocity field, we also consider the two eroded versions of sample **S1G0**, namely **S1G1** and **S1G2** (cf. Table 1). Visualisations of the simulation results for an inlet pressure of 2.1 bars are shown in Fig. 5(A). They qualitatively agree with the experimental findings of Proctor *et al*. [8], who provoked porous-permeable flows in meniscus samples through consolidation experiments at 2 bars. All cases in Fig. 5(A) have the velocity magnitudes normalised by the same maximum velocity obtained in **S1G0**. The effect of pore size on fluid flow is clearly visible.

In **S1G2**, the normalised velocity prediction is 120% higher than in **S1G0**, whereas in **S1G1**, the normalised velocity is only 30% higher than in **S1G0**. Using the erosion filter, the porosity increased by 20.5% in **S1G1** and 33.5% **S1G2**. Therefore, it is evident that the porosity of the meniscus medium has a significant and non-linear impact on fluid flow. This finding emphasises the importance of understanding the relationship between porosity and velocity in fluid flow.

The nonlinear flow characteristics can influence how effectively the meniscus distributes loads and lubricates the knee joint. Understanding these nonlinearities can provide insights into optimal joint function and highlight potential issues in cases of injury or degeneration.

## 3 Discussion

Understanding the impact of the microstructure of the human meniscus on its mechanical function is crucial for advancing our knowledge of knee joint biomechanics. Connecting the microstructural properties of the meniscus to its macroscopic function is still an unsolved problem, and investigating the impact of porosity and connectivity of the porous network within the meniscus on macroscopic fluid-mechanical properties can provide insights into physiological mechanisms. We therefore reconstructed the microstructure of two human menisci using high-resolution *µ*-CT scans (6,25*µ*m). By computational fluid dynamics (CFD) flow simulations with mesh-free particle methods and implicit boundaries, we show that the absorption properties of the meniscus are mainly explained by the diameters of the channels. For instance, a 30% increase in porosity leads to a 120% increase in velocity magnitude. These findings support the experimentally observed fact that the degraded function of aged menisci is associated with a condensed collagen network. Our findings contribute to further the understanding of the biomechanics of the meniscus and may have implications for the treatment of knee joint injuries and aging.

We used two *µ*CT scans (6.25 *µ*m resolution) of human meniscus samples [10] to study the effect of microstructure on meniscus fluid dynamics. We developed an end-to-end computational pipeline from *µ*CT scans to computational simulation results.

This was then used to simulate pore-scale flow using a mesh-free simulation method with different inlet pressure conditions.

For healthy sample geometries under physiological conditions, our simulations were able to reproduce two regimes of flow in accordance with the biophysical literature [6, 8, 11]: a visco-elastic creeping flow below a critical pressure of about 1.7 bar, and laminar porous-permeable flow above the threshold pressure. Interestingly, this threshold behavior was lost in both degraded sample geometries and at lower synovial fluid dynamic viscosity, both symptomatic of diseased states of the meniscus.

Fully resolved computer simulations of healthy menisci allowed us to understand the effect of the inlet pressure on flow patterns. This validated the method by comparison with experimental findings [6, 8, 11]. We then modified the healthy sample geometries outside the physiological range to study the effect of degeneration of the meniscus wall. This was achieved by way of an erosion filter used to explore several pore size configurations ranging from 0.53*µ*m to 0.80*µ*m at constantly high network connectivity. The resulting artificially eroded meniscus geometries were comparable to those seen from collagen fibers condensing in aging menisci [23, 38]. The simulation results showed that degenerating the meniscus wall non-linearly increases the transport velocity (+120% when increasing porosity by +33%) and decreases the meniscus’ tortuosity. The former impairs the meniscus’ functionality as a shock absorber, whereas the latter could hamper biochemical exchange in the tissue. According to our co-author Prof. MD. Seil, an expert in knee surgery, the erosion process identified in our research offers a potential translation for the reduction in cartilage and menisci thickness observed in MRI studies of individuals who participate in long-distance running [39]. This phenomenon is often associated with repetitive impact loading, especially in runners with intact menisci.

Our results showed that the pore size is critical in determining the fluid flow behavior, as smaller pores lead to capillary action slowing the flow, leading to a more uniform velocity distribution across pore sizes. Larger pores, on the other hand, reduce the fluid-solid interfacial area, resulting in reduced frictional forces and viscous drag, which leads to an increase in fluid velocity [40, 41]. However, the exact behavior can be influenced by the nature of the fluid (e.g., synovial fluid’s viscosity) and the specific microstructure of the porous media

We were also able to simulate the effect of aseptic loosening after total joint replacement [11], which provokes a decrease in synovial fluid viscosity. Indeed, the numerical results show that, if the dynamic viscosity of the synovial fluid is below physiological range, the threshold behavior of the interstitial flow, *i*.*e*. visco-elastic *vs*. porous-permeable, is lost. If this finding is confirmed by experiments, this could provide new avenues for therapy by modulating the viscosity back into the healthy range. Therefore, this dual behavior of the meniscus seems to be contained by the porous medium as the whole: the geometry of the microstructure coupled with the physiological range of synovial fluid dynamic viscosity.

Several leads for future improvement may be considered. First, experimental techniques have limitations we can not exclude the existence of artificial porosity provoked by freeze-drying protocol. However, the presented samples **S1G0** and **S2G0** show porosities within the physiological range of the knee meniscus [34], please refer to Table. 1. The present study reproduced the biophysical behavior of the human meniscus on only two samples. Reproduction of these results on a larger cohort is critical. Second, meniscus is considered in this study as a porous system composed of a rigid solid scaffold (*i.e*. a network of collagen fibers is not deformed) perfused by a viscous fluid. The present methodology could be used to confirm the hypothesis by Fithian *et al*. [6] that the deformation of the extra-cellular matrix may provoke porous-permeable flow. In order to reproduce this phenomenon, our mathematical model would need to be extended to consider poro-elastic effects [42], which are likely to play a key role there. In the present work, we also assumed that the pores are empty due to prior freeze-drying of the samples. We also did not model details of the solid scaffold of the meniscal tissue, such as the collagen network. Our model therefore did not include additional friction forces, but studied the role of microstructure geometry in isolation. Finally, the present simulations only considered static meniscus geometries. They did therefore not allow us to study pore collapse or recovery of the tissue from load, for which a simulation in a deforming geometry would be required. Future work could extend the algorithm in this direction, albeit this is not trivial.

To conclude, analysing fluid flow behavior in different realistic meniscus geometries using a direct numerical simulation method on a parallel multi-core computer allowed us to gain insight into how changes in porosity, connectivity, pore diameter, and fluid viscosity affect the macroscopic functional properties of human menisci. Our numerical simulations indicate that the observed threshold behavior of the meniscus as a porous medium is the result of a precise balance provided by the geometry of the microstructure and the dynamic viscosity of the synovial fluid. In the future, this knowledge and the presented numerical simulation program can be used to design better prosthetics and rehabilitation protocols, offering more effective and personalised treatment options.

## 4 Mathematical modelling

### 4.1 Entropically Damped Artificial Compressibility (EDAC)

In the realm of simulating the incompressible Navier-Stokes equations, Clausen [43]. introduced the EDAC method. This method paves the way for the explicit simulation of these equations. Within the EDAC formulation, a new equation governing the evolution of pressure p is introduced. This equation is derived from the thermodynamics of the system while maintaining a fixed density ρ. Remarkably, the EDAC method exhibits convergence to the incompressible Navier-Stokes (INS) equations when operating at low Mach numbers, and it maintains consistency both at low and high Reynolds numbers. Consequently, it becomes feasible to explicitly solve both the momentum equation and the pressure evolution equation within the Eulerian frame of reference,

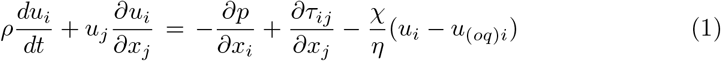

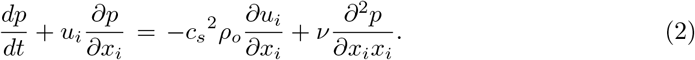

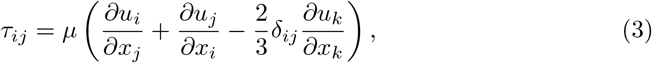

this approach involves the implicit penalisation of the computational domain, achieved by employing an indicator function χ that identifies the areas occupied by the solid geometry denoted as *O*.

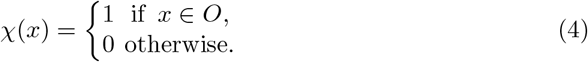

where, *u* is the velocity vector field, *p* is the pressure field, *t* is time, τ is the shear stress, *u*_(*oq*)*i*_ is the velocity of the solid body, *p*_(*oq*)*i*_ is the pressure in the solid body, ϕ is the porosity, and η = αϕ is the normalised viscous permeability. Note that 0 < ϕ ≪ 1 and 0 < η≪1, *µ* is the dynamic viscosity, χ is the penalisation mask function and *c*_*s*_ is the speed of sound.

The EDAC method converges to the incompressible Navier Stokes equations at low Mach numbers, and the equations can be solved explicitly in a Lagrangian or Eulerian frame of reference, please refer to [43].

### 4.2 Discretisation-Corrected Particle Strength Exchange (DC-PSE)

Discretisation-Corrected Particle Strength Exchange (DC-PSE) is a numerical method for consistently discretising differential operators on Eulerian or Lagrangian particles [44]. It is based on the following mollification or approximation of a sufficiently smooth function 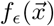 with a kernel η(),

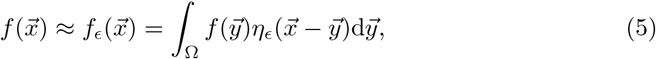

where ϵ is the length of the kernel for the supported particles. and the kernel is defined as

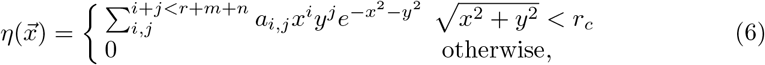

where the polynomial coefficients *a*_*i,j*_ are determined from discrete moment conditions evaluated at runtime over the given set of particles. The differential operators are derived using Taylor series expansion such that the operators are consistent for a desired order of convergence. We encourage the reader to refer to. [32] for further discussion regarding the EDAC formulation with the DC-PSE method, where the method is validated with different benchmarks and a convergence and accuracy study is provided.

### 4.3 Simulation setup, initial condition and boundaries

The DC-PSE simulations are performed in the Eulerian frame of reference with low Reynolds number *Re* ⪡ 1. The simulations use DC-PSE operators of convergence order 3, a particle interaction cutoff radius of 3.1ϵ, and second-order Adams Bashforth-Moulton adaptive time integration.

The boundary conditions are chosen according to the literature. At the inflow, we apply inlet pressure boundary conditions in *z* directions with a range of values (1.1 bar to 2.5) based on Refs. [6] and [8]. Periodic boundary conditions are applied in the *x* and *y* directions, while outflow conditions are implemented in the *z* direction. In the literature, various estimates for the fluid-dynamic viscosity *µ* have been provided [45, 46]. Bera *et al*. [46] found that there is no established standard for the dynamic viscosity of the synovial fluid, with a range of values from 0.7·10^*−*3^ to 3.5·10^*−*3^ Pa·s. Meanwhile, Fu *et al*. [45] determined that the viscosity of the synovial fluid in the meniscus with periprosthetic joint infection can be as high as 1.5·10^*−*2^ Pa ·s. Galandakova *et al*. [11] conducted a study to determine the synovial fluid viscosity in the knee joint using a Vibro viscometer and concluded the median viscosity value for a sample of 22 healthy menisci is 7.3·10^*−*3^ Pa · s. We use this viscosity value as a reference in our study.

The spatial domain is discretised using particles, with (128×128×256) particles in the *x, y*, and *z* directions, respectively. The DC-PSE solver for EDAC formulation is integrated into the C++ open-source high-performance computing platform OpenFPM. [47], (see Mathematical modelling). The C++ code is compiled using gcc 8.3.0 and OpenMPI 3.1.3 on 64-core AMD EPYC 7742 Processor (64 MB cache, 2.25 GHz) with 512GB RAM running Debian Linux 11.5.Each simulation time step takes approximately 0.4 seconds of wall-clock time when run on in parallel on all cores of a 64-core AMD EPYC 7742 processor.

### 4.4 From sample to computational domain

The STL geometry has surfaces that represent the walls of the meniscus, but these surfaces cannot be directly used in most numerical methods. To overcome this, we use an algorithm that labels the particles with a mask field χ [31] as fluid or solid depending on their position with respect to the STL surface. To study the mechanical behavior of the meniscus with varying porosity, the original geometry **S1G0** is taken as a reference, and modified geometries are generated with different volume fractions while conserving the connectivity of the porous network through a homogeneous erosion process. This process involves STL-to-voxel conversion, morphological erosion of the stack of voxels, voxel-to-STL conversion, and decimation and formatting to reduce computation cost.

## Acknowledgments

This work is funded by the Luxembourg National Research Fund (FNR) with the Core Junior grant lead by AO, “A Numerical homogenisation framework for characterising transport properties in stochastic porous media” (Por-Sol C20/MS/14610324). SPAB and SU acknowledge funding from a Luxembourg National Research Fund (FNR) grant number INTER/ANR/21/16399490.

## Author contributions

Author contributions: C.A.: Methodology, Software, Visualisation. S.U.: Investigation, Writing – original draft. I.F.S: Methodology, Review & Editing, Supervision. S.P.A.B: Methodology, Review & Editing. A.O: Conceptualisation, Methodology, Software, Investigation, Writing – original draft.

